# Proteomics of Extracellular Vesicles Produced by *Granulicatella* Species that cause Infective Endocarditis

**DOI:** 10.1101/2019.12.27.889220

**Authors:** Sarah A Alkandari, Radhika G Bhardwaj, Arjuna Ellepola, Maribasappa Karched

**Author notes:** Corresponding author Maribasappa Karched Oral Microbiology Research Laboratory Department of Bioclinical Sciences Faculty of Dentistry Kuwait University PO Box 24923 Safat 13110 Kuwait.

## Abstract

When oral bacteria accidentally enter the bloodstream due to transient tissue damage during dental procedures, they have the potential to attach to the endocardium or an equivalent surface of an indwelling prosthesis and cause infection. Many bacterial species produce extracellular vesicles (EVs) as part of normal physiology, but also use it as a virulence strategy. In this study, it was hypothesized that *Granulicatella* species produce EVs that possibly help them in virulence. Therefore, the objectives were to isolate and characterize EVs produced by these species and to investigate their immune-stimulatory effects. The reference strains *G. adiacens* CCUG 27809 and *G. elegans* CCUG 38949 were cultured on chocolate blood agar for 2 days. From subsequent broth cultures, the EVs were isolated using differential centrifugation and filtration protocol and then observed using scanning electron microscopy. Proteins in the vesicle preparations were identified by nano LC-ESI-MS/MS. The EVs proteomes were analyzed and characterized using different bioinformatics tools. The immune-stimulatory effect of the EVs was studied via ELISA quantification of IL-8, IL-1β and CCL5, major proinflammatory cytokines, produced from stimulated human PBMCs. It was revealed that both *G. adiacens* and *G. elegans* produced EVs, ranging in diameter from 30 to 250 nm. Overall, *G. adiacens* EVs contained 160 proteins, and *G. elegans* EVs contained 107 proteins. Both proteomes consist of several ribosomal proteins, DNA associated proteins, binding proteins, and metabolic enzymes. It was also shown that these EVs carry putative virulence factors including moonlighting proteins. These EVs were able to induce the production of IL-8, IL-1β and CCL5 from human PBMCs. The diversity in EVs content indicates that these vesicles could have possible roles in bacterial survival, invasion, host immune modulation as well as infection. Further functional characterization of the *Granulicatella* EVs may provide new insights into virulence mechanisms of these important but less studied oral bacterial species.

## Introduction

*Granulicatella* species, formerly known as nutritionally variant streptococci based on their characteristic dependence on pyridoxal or cysteine supplementation for their growth in standard media [1], are catalase and oxidase negative, non-motile, non-spore-forming, facultatively anaerobic Gram-positive cocci [2, 3]. They are part of the normal oral flora [4], but cause serious infections such as infective endocarditis. The genus *Granulicatella* consists of 3 species: *Granulicatella adiacens*, *Granulicatella elegans* and *Granulicatella balaenopterae* [3]. The species *G. balaenopterae* has not been isolated from human samples, whereas both *G. adiacens* and *G. elegans* have been reported from IE cases [5, 6]. In addition, these oral commensal cocci have been associated with endodontic infections [7, 8], dental caries [9], and periodontitis [8, 10] via DNA-based studies. Although this association does not substantiate the role of *Granulicatella* species in dental diseases, the fact that these species are causative agents in infective endocarditis implies that they might exert similar pathogenic potential also in the oral cavity.

Many bacterial species routinely produce extracellular vesicles (EVs) during normal growth [11]. Gram-negative bacteria are commonly found to produce such vesicles, which are derived from blebbing of the outer membrane and thus are called outer membrane vesicles (OMVs) [11]. Generally, these OMVs contain outer membrane proteins, lipopolysaccharides, glycerophospholipids in addition to enclosed periplasmic components and bacterial nucleic acids [11–13]. The study of the EVs was initially limited to Gram-negative bacteria, as it was thought that the rigidity of the Gram-positive cell wall, which is rich in peptidoglycans, would not allow vesicle blebbing [11]. However, the production of EVs was also observed in some Gram-positive bacteria [14–17]. Current studies [18–20] showed that the activity of cell wall-degrading enzymes, which weaken the peptidoglycan layer and thus facilitate the release of Gram-positive EVs, could probably explain such phenomena in Gram-positive bacteria. Similar to Gram-negative OMVs, these EVs contain proteins, lipids, enzymes, toxins and bacterial nucleic acids [20]. However, Gram-positive EVs can still be distinguished from OMVs as the former lack lipopolysaccharide and enclosed periplasmic components [20].

Several studies [13, 14, 18] showed that bacteria exploit vesicle production as a virulence strategy. Bacterial components, including virulence factors, are packed in the vesicles and delivered to the host cells and tissues. The vesicle-derived virulence factors play an important role in bacterial pathogenicity, e.g., by eliciting an inflammatory response, manipulating the host’s immunity, eliminating the competing commensal microorganisms, relieving internal stress, mediating biofilm formation, and acting as decoys absorbing and blocking cell wall-lytic compounds and membrane-disrupting antimicrobial peptides produced by other commensals and host innate immune cells [13, 14, 18].

Protein secretion in *Granulicatella* species has been studied [21], but vesicle production in these species has not been investigated yet. In this study, it was hypothesized that *Granulicatella* species produce EVs that possibly play a role in the pathogenesis of *Granulicatella* infections.

## Materials and methods

### Bacterial strains and culture conditions

The reference strains *G. adiacens* CCUG 27809 and *G. elegans* CCUG 38949 were cultured on chocolate blood agar (CBA) with 0.001% pyridoxal hydrochloride at 37 °C and in 5% CO_2_ in air for 2 days. A loop-full of colonies from the CBA plates was inoculated into brucella broth supplemented with 0.001% pyridoxal hydrochloride and incubated as above for 2 days.

### Isolation of EVs

The EVs were isolated using a previously described centrifugation and filtration protocol [22], with slight modifications. Briefly, for pelleting the bacteria, the broth culture was centrifuged at 5000 × g at room temperature for 10 minutes (Centrifuge 5430 R, Eppendorf AG, Germany). For removing any remnants of intact bacterial cells, the supernatant was filtered through a 0.22 µm sterile syringe filter (Millipore, Germany). The filtrate was then re-centrifuged at 125000 × g at 4° C for 3 hours (Optima™ L-XP ultracentrifuge, Beckman, USA). The obtained pellet was suspended in 300 µl sterile phosphate-buffered saline (PBS). The EVs samples were stored at -20° C until used.

### Preparation of whole cell protein (WCP)

A loop full of colonies from the CBA plates was suspended in 2 ml sterile PBS. The bacterial suspension was centrifuged at 5000 × g at room temperature for 5 minutes (Centrifuge 5430 R, Eppendorf AG, Germany). Then, after discarding the supernatant, the pellet was washed with 2 ml sterile PBS. The bacterial whole cell protein (WCP) was obtained by ultra-sonicating bacterial cells at 40 pulse rate on ice for 8 cycles (1 minute sonication followed by 1 minute rest per cycle) (Omni Sonic Ruptor 4000, Omni International, USA) followed by centrifugation at 7000 × g at 4° C for 10 minutes (Centrifuge 5430 R, Eppendorf AG, Germany). The resulting supernatant was used as the WCP sample and stored at -20° C until used.

### Characterization of EVs

#### Scanning electron microscopy (SEM)

The obtained vesicle preparations were suspended in sterile PBS containing 3% glutaraldehyde for 2 hours on a rotator and then kept in a refrigerator overnight. For staining, the vesicle samples were incubated in 1% osmium tetroxide for 2 hours. For dehydration, the samples were kept in increasing concentrations of acetone from 30 to 100%, 10 minutes in each, on a rotator. The samples were then placed in a critical point dryer for complete drying, mounted on stubs with carbon double adhesive tape and finally coated with gold and stored in a desiccator until observation. The samples were observed on Zeiss Leo Supra 50VP field emission scanning electron microscope (Carl Zeiss, Germany). For comparison, SEM analysis of bacterial whole cells was also performed using the same previous biological sample preparation protocol.

#### Determination of protein concentration and SDS-PAGE

Protein concentrations in the EVs and WCP samples were determined by Quick Start^TM^ Bradford protein microplate standard assay (Bio-Rad, USA). For protein separation, the samples were subjected to sodium dodecyl sulfate-polyacrylamide gel electrophoresis (SDS-PAGE) using the mini-PROTEAN II cell electrophoresis system (Bio-Rad, USA). The proteins were denatured in 2× loading buffer at 100°C for 5 minutes, followed by centrifugation at 5000 ×g for 5 minutes. 20 µl of proteins loaded in each well of the gel were separated on 12% SDS-PAGE at a constant 120 V. After the run was completed, protein bands were detected using silver stain. Gel images were visualized in G: Box Imaging System (Syngene, India). Protein banding patterns and molecular weights of the bands were determined using GeneSys tools software

#### Identification of EVs proteins by Nano-LC-ESI-MS/MS

For the identification of EVs proteins, mass spectrometry was performed by Proteome Factory (Proteome Factory AG, Berlin, Germany) using nano-liquid chromatography-electrospray ionization-tandem mass spectrometry (nano-LC-ESI-MS/MS). After pooling replicate samples from EVs preparations, 400 ng proteins were reduced, alkylated and digested by trypsin (Promega, Mannheim, Germany). Then, the resulting peptides were subjected to the nanoLC-ESI-MS/MS. 1% acetonitrile/0.5% formic acid was used as eluent for 5 minutes to trap and desalt the peptides on the enrichment column (Zorbax SB C18, 0.3 × 5 mm, Agilent). An acetonitrile/0.1% formic acid gradient from 5% to 40% acetonitrile was then used within 120 minutes to separate the peptides on a Zorbax 300 SB C18, 75 µm x 150 mm column (Agilent). The mass spectrometer automatically recorded the mass spectra according to the manufacturer’s settings. Protein identification was made using the Mascot search engine (Matrix Science, London, England) and the National Center for Biotechnology Information non-redundant (NCBI-nr) protein database, version 20151202, (NCBI, Bethesda, USA). Ion charge in search parameters for ions from ESI-MS/MS data acquisition was set to “1+, 2+ or 3+” according to the instrument’s and method’s standard charge state distribution. The search parameters were: Fixed modifications: Carbamidomethyl (C); variable modifications: Deamidated (NQ), Oxidation (M); Peptide Mass Tolerance: ± 5 ppm; Fragment Mass Tolerance: ± 0.6 Da; Missed Cleavages: 2. The inclusion criterion was: peptides that match with a score of 20 or above. Mass spectrometry data, with the project acession number PXD015630, has been deposited at PRIDE archive (https://www.ebi.ac.uk/pride/archive/) repository. The data files can be accessed with the username and the password **k0SBY6YK**.

#### Bioinformatic analysis

Protein sequences from the liquid chromatography-mass spectrometer (LC-MS) analysis of the EVs proteomes were analyzed by an *in silico* 2-dimensional electrophoresis (2-DE) tool. For this, the software JVirGel, version 2.0 (http://www.jvirgel.de/index.html), was used to obtain a theoretical (2-DE) image of the EVs proteins [23]. The subcellular localization of the EVs proteins detected with LC-MS/MS was predicted using the PSORTb tool, version 3.0.2 (https://www.psort.org/psortb/) [24]. To determine if any of the secreted proteins are packed into the vesicles, the prediction tool SignalP, version 5.0 (http://www.cbs.dtu.dk/services/SignalP/abstract.php), was utilized to predict proteins secreted via the general *Sec*retion route (Sec-pathway) [25]. In addition to that, the prediction tool TatP (http://www.cbs.dtu.dk/services/TatP/), was used to predict proteins secreted via the Twin-arginine translocation pathway (Tat-pathway) [26]. To identify lipoproteins, lipoboxes were searched using the prediction tools LipoP (http://www.cbs.dtu.dk/services/LipoP/) and PRED-LIPO (http://bioinformatics.biol.uoa.gr/PRED-LIPO/input.jsp) [27].

#### Function prediction analysis

Proteins with multiple functions, known as “moonlighting proteins”, were identified using the prediction tool moonprot, version 2.0 (http://www.moonlightingproteins.org/) [28], and searching the database Multitask ProtDB (http://wallace.uab.es/multitaskII/) [29]. Gene Ontology (GO) analysis of the EVs proteomes was performed using the amino acid FASTA sequences of *G. adiacens* and *G. elegans*. For this, GO annotations were analyzed and plotted using the tools Blast2GO (https://www.blast2go.com/) [30], and WEGO, version 2.0 (http://wego.genomics.org.cn/) [31]. The EVs proteins were grouped based on functional association networks using the tool STRING (https://string-db.org/) [32].

Minimum interaction scores were set at a strong confidence level of 0.7. The EVs proteins were also grouped based on different biological pathways. For this, all protein sequences from *G. adiacens* and *G. elegans* EVs proteomes were analyzed by the Kyoto Encyclopedia of Genes and Genome (KEGG) (https://www.genome.jp/kegg/pathway.html) pathway analysis tool using the genus “streptococcus” as reference [33].

#### Prediction of virulence factors in the EVs proteomes

To predict virulence proteins in the EVs proteomes, the tool VirulentPred (http://203.92.44.117/virulent/) [34], along with the Virulence Factor Data Base (VFDB; http://www.mgc.ac.cn/VFs/) were used. Proteins predicted to be virulent by the previous tools were manually searched in the literature for experimental evidence on their virulence properties.

### Cytokine induction of human PBMCs by EVs

#### Isolation of human PBMCs

PBMCs from the blood of a healthy human volunteer were isolated using Ficoll-Paque density gradient centrifugation method [35]. After obtaining written informed consent from the donor, blood was collected by venipuncture into vacutainer heparin tubes (3 ml per tube). The blood was then carefully layered onto 3.5 ml Ficoll-Paque media solution (GE Healthcare, USA) in a sterile centrifugation tube. For separating mononuclear cells, the tubes were centrifuged at 3400 ×g at room temperature with the brakes off for 10 minutes. The layer of PBMCs, the buffy coat layer, was then transferred to another sterile centrifugation tube. The cell isolate was washed twice by resuspending it in 5 ml RPMI medium followed by centrifugation at 2000 rpm at room temperature with the brakes on for 5 minutes. The supernatant was discarded, and the cell pellet was finally resuspended in 1 ml RPMI medium supplemented with 10% heat-inactivated fetal bovine serum and 2% Gibco^TM^ 100× antibiotic-antimycotic solution. Cell concentration in the PBMCs sample was estimated by loading 10 µl aliquot on a hemocytometer under 400× magnification.

#### Stimulation of human PBMCs with EVs and WCP

Isolated human PBMCs were stimulated with different concentrations (10, 25, 50, and 100 µg/ml) of *G. adiacens* EVs, *G. adiacens* WCP, *G. elegans* EVs, and *G. elegans* WCP for 24 hours. For this, in a 24-well plate, 480 µl supplemented RPMI medium containing PBMCs (10^6^ cells per ml) was added to each well and stimulated with 20 µl of bacterial EVs or WCP. The plate was incubated at 37 °C and in 5% CO_2_ in air for 24 hours. Well with 20 µl sterile PBS and 480 µl RPMI medium containing PBMCs was used as negative control.

#### Quantitative determination of selected cytokines

The quantitative sandwich enzyme-linked immunosorbent assay (ELISA) technique was used to quantify the production of the human cytokines IL-8, IL-1β, and CCL5 (RANTES) from the stimulated PBMCs. For this, ELISA immunoassay kits (Quantikine^®^ ELISA R&D systems, Bio-Techne, USA) were used according to the manufacturer’s instructions. Briefly, standards, samples, and controls were added to the wells of a 96-well microplate pre-coated with a monoclonal antibody specific for the cytokine of interest. To allow the specific cytokine in the sample to be bound by the specific immobilized antibody, the plate was incubated at room temperature for 2 hours. To remove any unbound substances, the wells were washed with wash buffer using ImmunoWash^TM^ 1575 microplate washer (Bio-Rad, USA). Then, an enzyme-linked polyclonal antibody for the specific cytokine was added to each well. After an incubation period of one hour at room temperature, the wells were washed again with wash buffer to remove any unbound antibody-enzyme reagent. A substrate solution was then added to each well, and the microplate was incubated at room temperature for 20-30 minutes while being protected from light. To terminate the colorful enzyme-substrate reaction, a stop solution was added to each well. Finally, iMark^TM^ microplate reader was used to measure the intensity of the color developed.

### Statistical analysis

All experiments were repeated twice. Statistical Package for Social Sciences Software (SPSS), version 25, was used for data analysis. Descriptive statistics were presented using mean ± standard deviation (SD). Independent-samples T test and Mann Whitney U test were used to analyze differences between groups. A critical probability value (P value) of < 0.05 was used as the cut-off level for statistical significance.

### Ethical considerations

This study was approved by the ethical committee of the Health Sciences Center, Kuwait University (DR/EC/3413), and has been carried out in full accordance with the World Medical Association Declaration of Helsinki. The blood donor received written information about the nature and purposes of the study and a written informed consent was obtained upon his/her approval to participate.

## Results and discussion

### Isolation of EVs

It was revealed by the current study that both *G. adiacens* and *G. elegans* produce EVs. Vesicles of varying sizes, ranging from 30 to 250 nm in diameter, were seen in the electron micrographs. This nano-scale range size was consistent with other bacterial EVs [14, 16]. For comparison, images of bacterial whole cells (Figs 1A and 2A) and the vesicle preparations (Figs 1B and 2B) were captured at the same magnification of ×10000. Vesicle shape and size could be visualized better at a higher magnification of × 40000 (Figs 1C and 2C).

**Fig 1.**
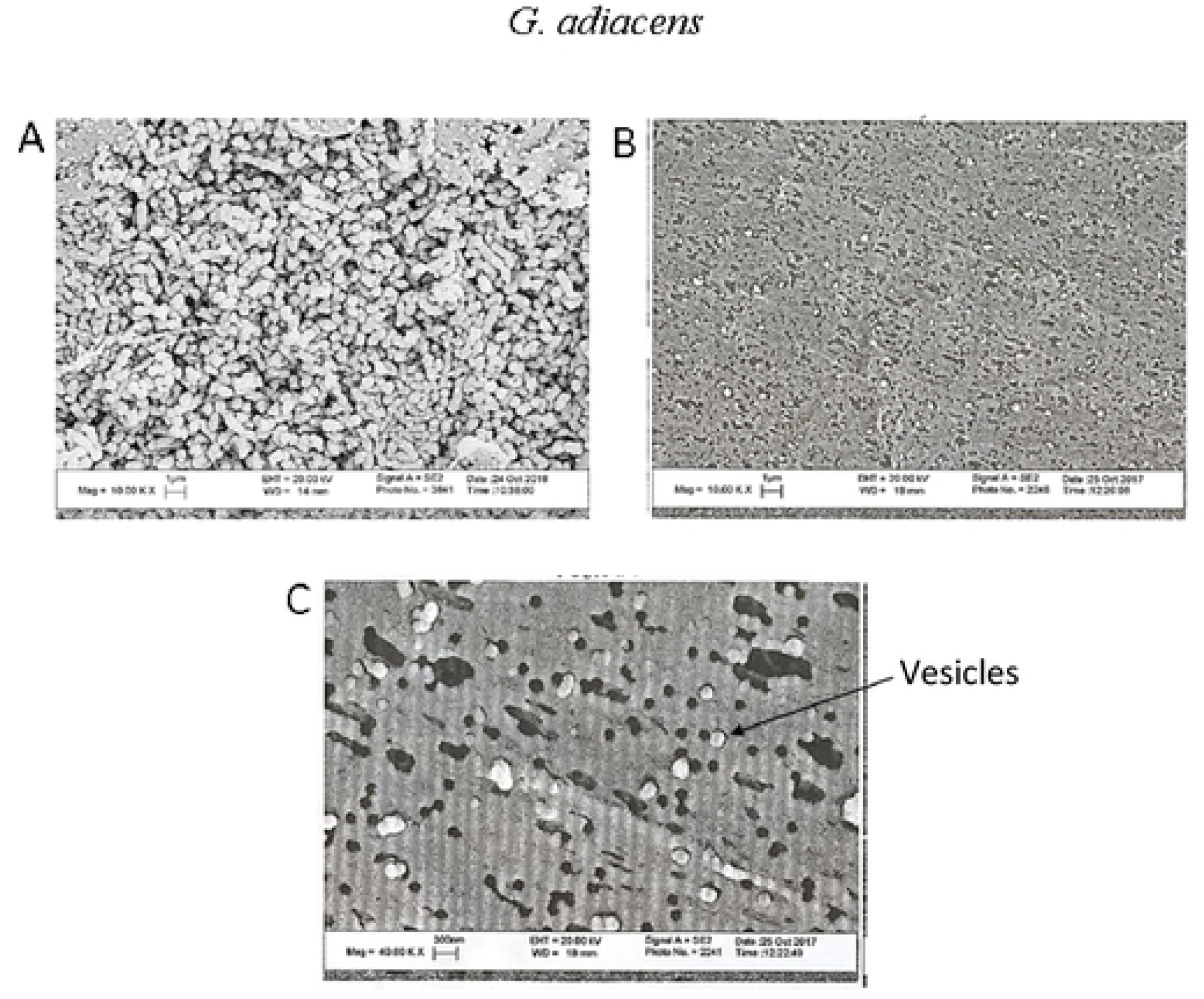
SEM images of *G. adiacens* whole cells and the EVs preparation. SEM images of bacterial whole cells (A) and the EVs preparation (B) captured at the magnification ×10000. (C) SEM images of the EVs acquired at × 40000.

**Fig 2.**
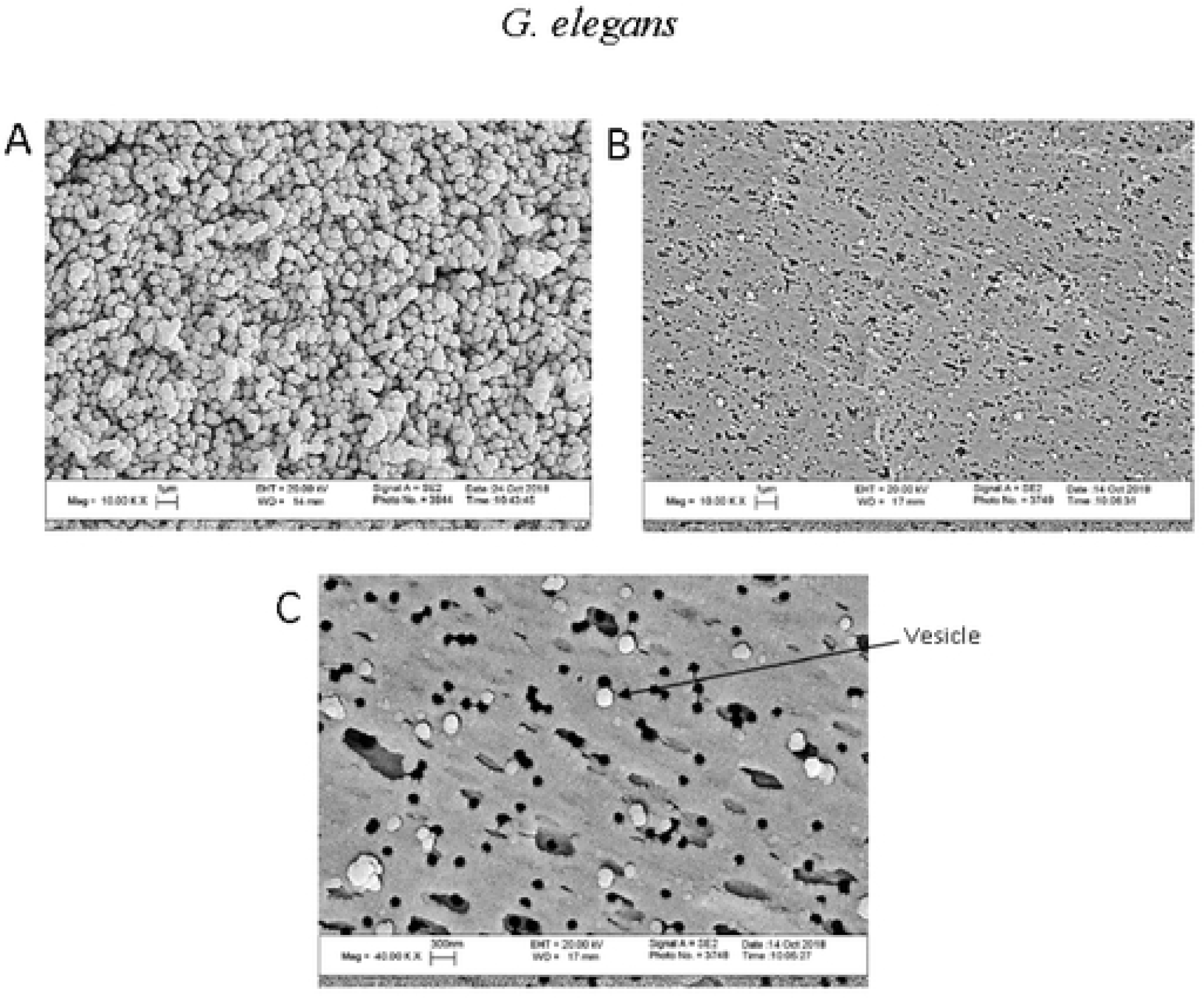
SEM images of *G. elegans* whole cells and the EVs preparation. SEM images of bacterial whole cells (A) and the EVs preparation (B) captured at the magnification ×10000. (C) SEM images of the EVs acquired at × 40000.

### Characterization of EVs

#### Determination of protein concentration and SDS-PAGE

Protein concentrations in the EVs samples from both *G. adiacens* and *G. elegans*, 1337 µg/ml and 1339 µg/ml respectively, were much lower compared to their respective WCP samples, 3102 µg/ml and 3388 µg/ml respectively. Consistently, SDS-PAGE analysis revealed that the EVs preparations from both *G. adiacens* and *G. elegans* showed much fewer bands on gel than their respective WCP preparations (Fig 3A).

**Figure 3.**
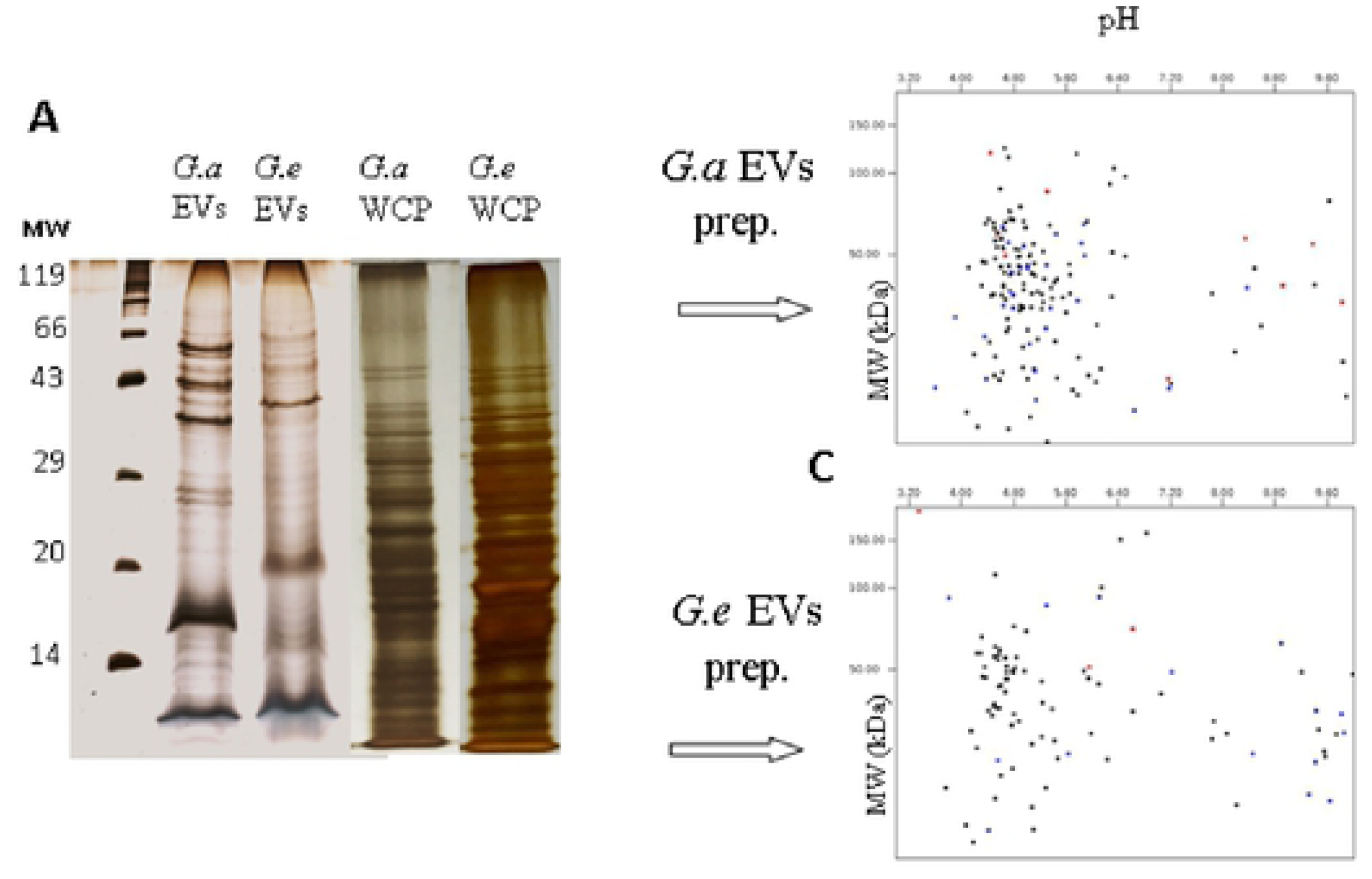
Analysis of the proteome of *G. adiacens* and *G. elegans* EVs. (A) SDS-PAGE gel showing protein bands from EVs and WCP preparations. (B) and (C) Protein sequences from LC-MS analysis of the vesicle proteome analyzed by an *in silico* 2-DE tool.

#### Identification of EVs proteins by NanoLC-ESI-MS/MS

In total, 160 and 107 proteins detected by NanoLC-ESI-MS/MS in EVs preparations of *G. adiacens* and *G. elegans* respectively, were analyzed and defined as the EVs proteomes in the present study (Suppl. Table 1 and 2). These numbers were within the range of proteins identified in previous analyses of other bacterial vesicle proteomes [14, 16, 17].

#### Bioinformatic analysis

*In silico* 2-DE analysis of the EVs proteomes showed that the molecular mass of the proteins ranged between 14.9 kDa and 125.7 kDa for *G. adiacens* (Fig 3B), and between 20 kDa and 195 kDa for *G. elegans* (Fig 3C). The proteome of *G. adiacens* EVs formed a distinct cluster with respect to predicted isoelectric point (pI) values in the range of 4.0 and 6.4 (Fig 3B). In the case of *G. elegans* EVs, the proteins seemed to be more dispersed in the pI range, showing some clustering of proteins in the pI range 4.0 to 5.6 (Fig 3C).

According to the PSORTb subcellular localization prediction tool analysis, *G. adiacens* EVs proteome was predicted to contain 113 cytoplasmic proteins, 27 cytoplasmic membrane proteins, and 4 cell-wall anchored proteins; whereas the localization of 16 proteins could not be predicted. Similarly, *G. elegans* EVs proteome was predicted to contain 67 cytoplasmic proteins, 20 cytoplasmic membrane proteins, 2 cell-wall anchored proteins and 18 proteins of unknown localization. As predicted in this study, the majority of EVs proteins were cytoplasmic in both *G. adiacens* and *G. elegans* proteomes (71% and 67% respectively). Cytoplasmic proteins located in other bacterial vesicles have been reported in several earlier studies [36, 38]. Existing evidence suggests that the enormous location of cytoplasmic proteins into vesicles is due to specific sorting mechanisms, and not due to lysis of dead cells [39]. Importantly, cytoplasmic proteins released as part of vesicles are known to function as adhesins, contribute to biofilm matrix formation, and help bacteria in evading the immune system [40].

As predicted in our study by the SignalP and TatP tools, secretory proteins were packed into the EVs of both *Granulicatella* species. According to the SignalP prediction tool, 38 proteins of *G. adiacens* EVs proteome were found to contain a signal sequence, while 18 of the *G. elegans* EVs proteins were predicted to contain a signal sequence. This suggests that such proteins are probably secreted via the Sec-pathway. The TatP prediction tool showed that 12 proteins of the *G. adiacens* EVs proteome and 13 proteins of the *G. elegans* EVs proteome contained TatP signal sequence, suggesting the Tat pathway for their secretion. Both the Sec and Tat pathways are major pathways that exist in bacteria for proteins secretion across the cytoplasmic membrane [41, 42]. The former pathway is well known to translocate proteins in their unfolded conformation, while the latter catalyzes the secretion of proteins that fold before their translocation [42]. It is well-established that protein secretion is an essential strategy in the pathogenesis of bacterial infections [41]. “Secreted proteins can play many roles in promoting bacterial virulence, from enhancing attachment to eukaryotic cells, to scavenging resources in an environmental niche, to directly intoxicating target cells and disrupting their functions” [41]. Lipoprotein prediction tools (Pred-Lipo, LipoP) revealed that there were 23 lipoproteins in the *G. adiacens* EVs proteome, and 10 lipoproteins in the *G. elegans* EVs proteome.

#### Function prediction analysis

The present study showed that EVs from both *Granulicatella* species carry proteins predicted to exhibit multitasking capabilities. Table 1 lists the 12 proteins from the *G. adiacens* EVs proteome and the 7 proteins from the *G. elegans* EVs proteome, respectively, that were identified as “moonlighting proteins”. Major proteins predicted as multifunctional proteins were ribosomal proteins and molecular chaperones. Additionally, a glycolytic enzyme, glyceraldehyde-3-phosphate-dehydrogenase, and a few putative virulent proteins such as NADH oxidase and thioredoxin were also identified. Such multifunctional bacterial proteins were found to play a role in the virulence of several other human pathogenic bacteria; e.g., *Staphylococcus aureus*, *Streptococcus pyogenes*, *Streptococcus pneumoniae*, *Helicobacter pylori*, and *Mycobacterium tuberculosis* [43–45].

**Table 1.** Predicted moonlighting proteins from G. adiacens and G. elegans EVs proteome.

| GI Number | Protein |
| --- | --- |
| **<u>*G. adiacens*</u>** |  |
| gi\|491802570 | Serine protease |
| gi\|491797953 | Molecular chaperone DnaK |
| gi\|491800441 | Superoxide dismutase |
| gi\|491800797 | Glyceraldehyde-3-phosphate dehydrogenase |
| gi\|491800365 | NADH oxidase |
| gi\|748591028 | 30S ribosomal protein S20 |
| gi\|491801600 | 50S ribosomal protein L7/L12 |
| gi\|491802592 | 30S ribosomal protein S6 |
| gi\|259036192 | Thioredoxin |
| gi\|491801148 | Elongation factor Tu |
| gi\|491801605 | 50S ribosomal protein L10 |
| gi\|259035990 | Phosphoglycerate kinase |
| **<u>*G. elegans*</u>** |  |
| gi\|491797953 | Molecular chaperone DnaK |
| gi\|491800797 | Glyceraldehyde-3-phosphate dehydrogenase |
| gi\|491800365 | NADH oxidase |
| gi\|491799730 | Short-chain dehydrogenase |
| gi\|491801600 | 50S ribosomal protein L7/L12 |
| gi\|491802592 | 30S ribosomal protein S6 |
| gi\|491801148 | Elongation factor Tu |

Fig 4 summarizes the Gene Ontology analysis of the EVs proteomes. Overall, 112 of the *G. adiacens* sequences and 108 of the *G. elegans* sequences were assigned with GO annotation. For *G. adiacens* and *G. elegans*, the proteins were divided into 3 groups based on GO terms: 90 and 61 proteins in “biological process” group, 28 proteins each in the “cellular component” group, and 104 and 70 proteins in the “molecular function” group, respectively. According to the Gene Ontology analysis conducted in the present study, most proteins in both *G. adiacens* EVs and *G. elegans* EVs proteomes were predicted to be involved in molecular functions, particularly catalytic and binding functions, followed by biological processes, mainly metabolic and cellular processes. It is possible that these species might utilize nutrients in the environment by using the metabolism-mediator proteins in the EVs [46]. Only 28 proteins in both proteomes were annotated for cellular components. Similar to other bacterial EVs, *G. adiacens* and *G. elegans* EVs contained several ribosomal proteins, DNA associated proteins, binding proteins, and metabolic enzymes, indicating that bacterial EVs might facilitate the transfer of functional proteins [14, 18].

**Fig 4.**
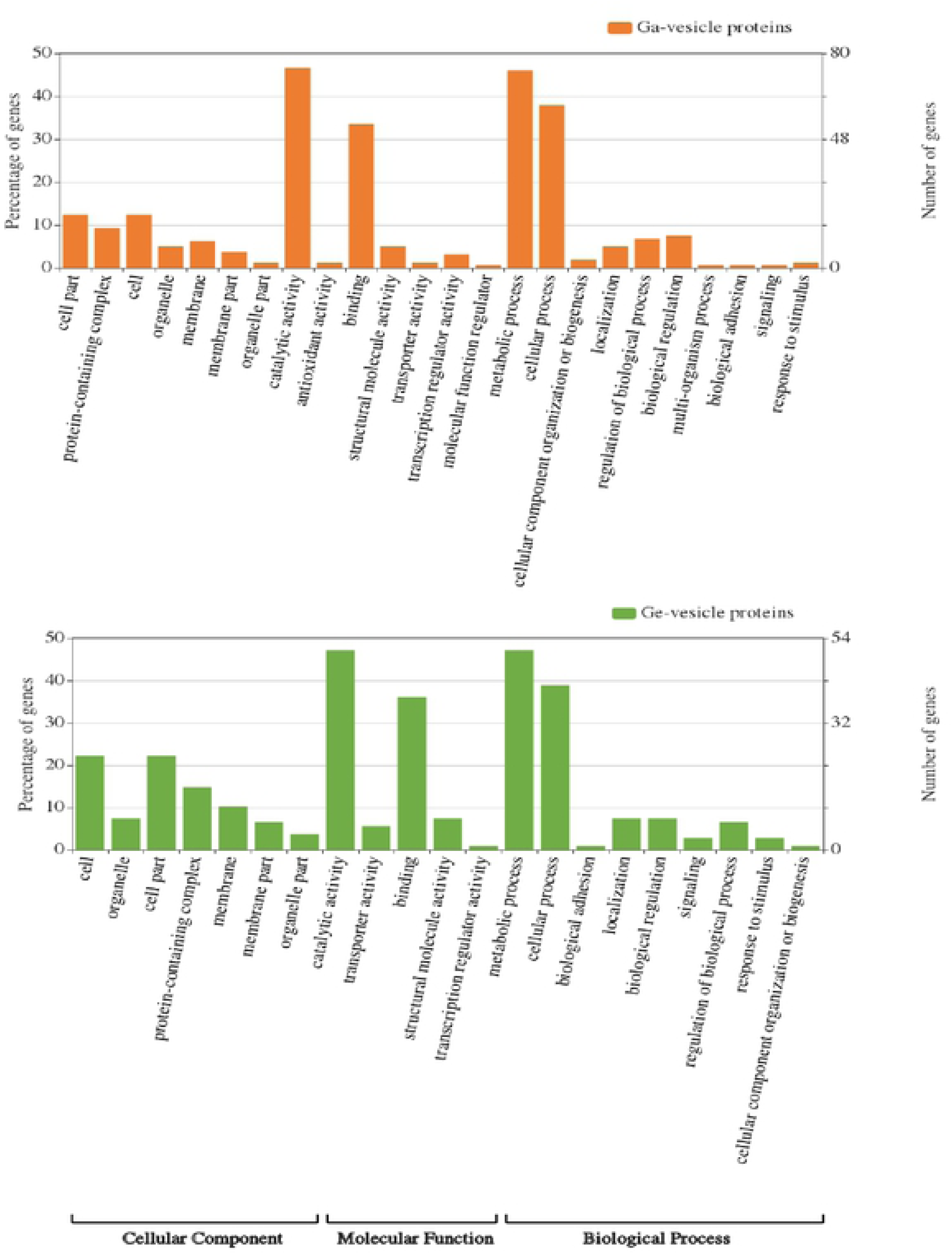
Gene Ontology analysis of the proteomes of *G. adiacens* and *G. elegans* EVs preparations. Gene ontology annotation was achieved using Blast2GO and an online software “WEGO”. Protein sequences were grouped into 3 categories based on their properties and functions.

Figures 5 and 6 demonstrate the STRING functional protein association network analysis of *G. adiacens* EVs proteome and *G. elegans* EVs proteome, respectively. As demonstrated in our study, both *G. adiacens* and *G. elegans* EVs proteomes formed three distinct protein groups based on their functional associations. These groups were carbohydrate metabolism, ribosomal proteins, and heat shock proteins/chaperones. Components of the carbohydrate metabolism network were: glyceraldehyde-3-phosphate dehydrogenase, phosphoenolpyruvate-protein phosphotransferase, glucose-6-phosphate isomerase, phosphoglycerate kinase, Pyruvate kinase, ATP-dependent 6-phosphofructokinase, transketolase, pyruvate dehydrogenase E1 component, and dihydrolipoamide acetyltransferase component of pyruvate dehydrogenase complex. The ribosomal protein group consisted mainly of the secreted ribosomal proteins: 30S ribosomal protein S20, 50S ribosomal protein L10, 30S ribosomal protein S5, 50S ribosomal protein L5, 50S ribosomal protein L7/L12, 30S ribosomal protein S6; Binds together with S18 to 16S ribosomal RNA, 50S ribosomal protein L11, 30S ribosomal protein S7, 50S ribosomal protein L2, Ribosome-recycling factor, and 50S ribosomal protein L1. Putative virulence-associated proteins, thioredoxin, superoxide dismutase and molecular chaperones (DnaK, DnaN, GroL, and GrpE) formed another cluster. In the case of *G. elegans*, DnaK was the only chaperone found. A growing body of literature [43–45] has shown that a number of enzymes involved in the glycolytic pathway as well as molecular chaperones are recognized as moonlighting proteins and thus could play a role in the pathogenesis of bacterial infection. Of the glycolytic enzymes detected in EVs proteomes in this study, glyceraldehyde-3-phosphate dehydrogenase, glucose-6-phosphate isomerase, phosphoglycerate kinase, pyruvate kinase, and ATP-dependent 6-phosphofructokinase were found to possess moonlighting properties. These enzymes could function as transferrin receptor, cell signaling kinase, neutrophil evasion protein, immunomodulator, plasminogen binding protein, fibrinogen binding protein, actin binding protein, and has a role in NAD-ribosylation activity and extracellular polysaccharide synthesis [44]. Moreover, the molecular chaperone DnaK was found to act as a multifunctional protein, which could stimulate CD8 lymphocyte and monocyte chemokines production, compete with HIV for binding to CCR5 receptors, and bind plasminogen [44]. In addition, it was concluded by a previous study [47] that many bacterial ribosomal proteins could function beyond their primary role as ribosomes, integral components of protein synthesis machinery. These proteins could also modulate different cell processes, such as transcription, regulation of the mRNA stability, DNA repair and replication, and phage RNA replication [47]. Furthermore, the L7/L12 ribosomal protein was experimentally proven to elicit a cell-mediated immune response in mice [48].

**Fig 5.**
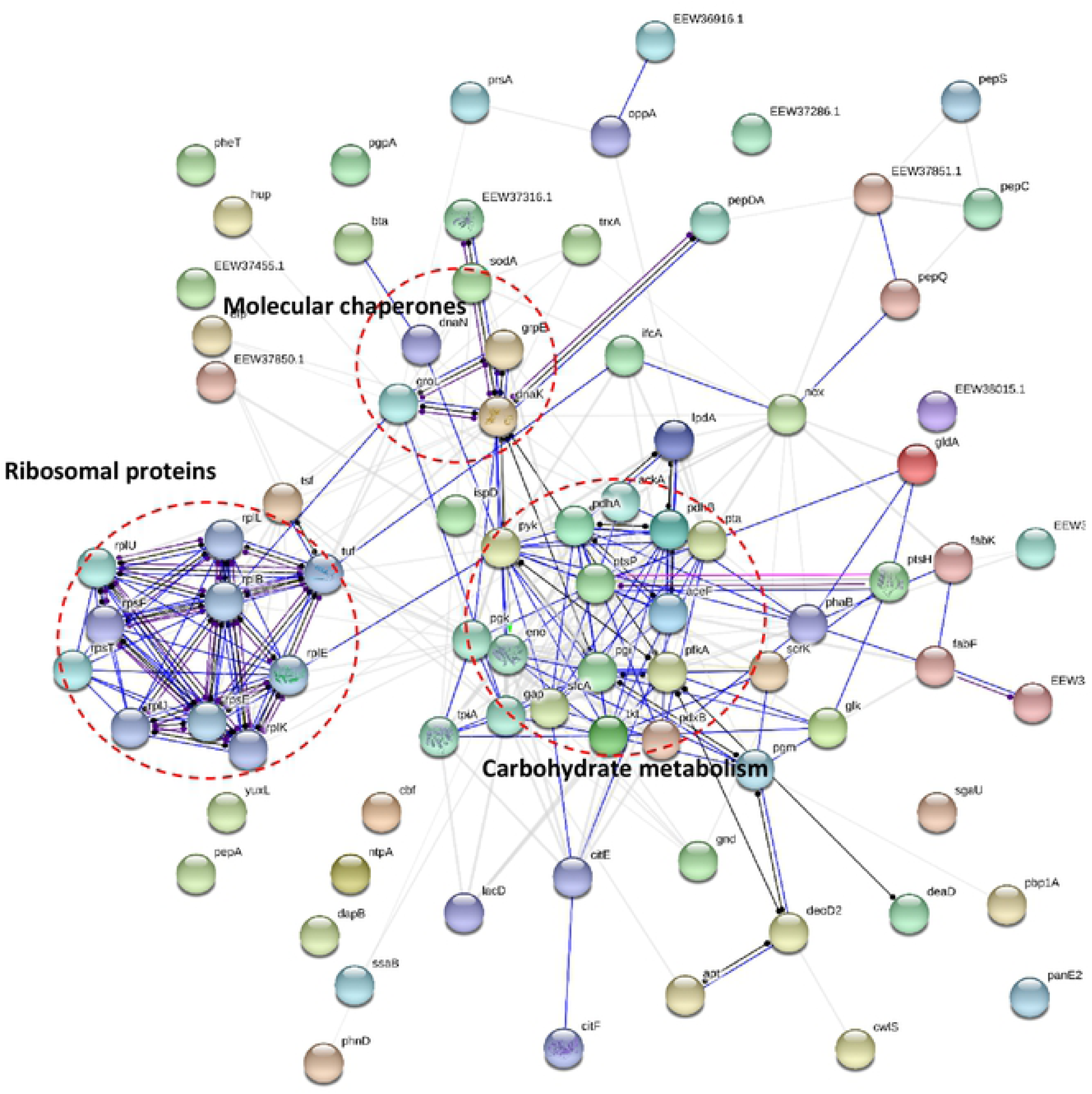
Functional protein association networks of *G. adiacens* EVs proteome. The online tool STRING was used for grouping the EVs proteins based on functional networks. Minimum interaction scores were set at a strong confidence level of 0.7. The three major network groups formed are shown in dotted circles. Seven different colors link a number of nodes and represent seven types of evidence used in predicting associations. A red line indicates the presence of fusion evidence; a green line represents neighborhood evidence; a blue line represents co-occurrence evidence; a purple line represents experimental evidence; a yellow line represents text mining evidence; a light blue line represents database evidence and a black line represents co-expression evidence.

**Fig 6.**
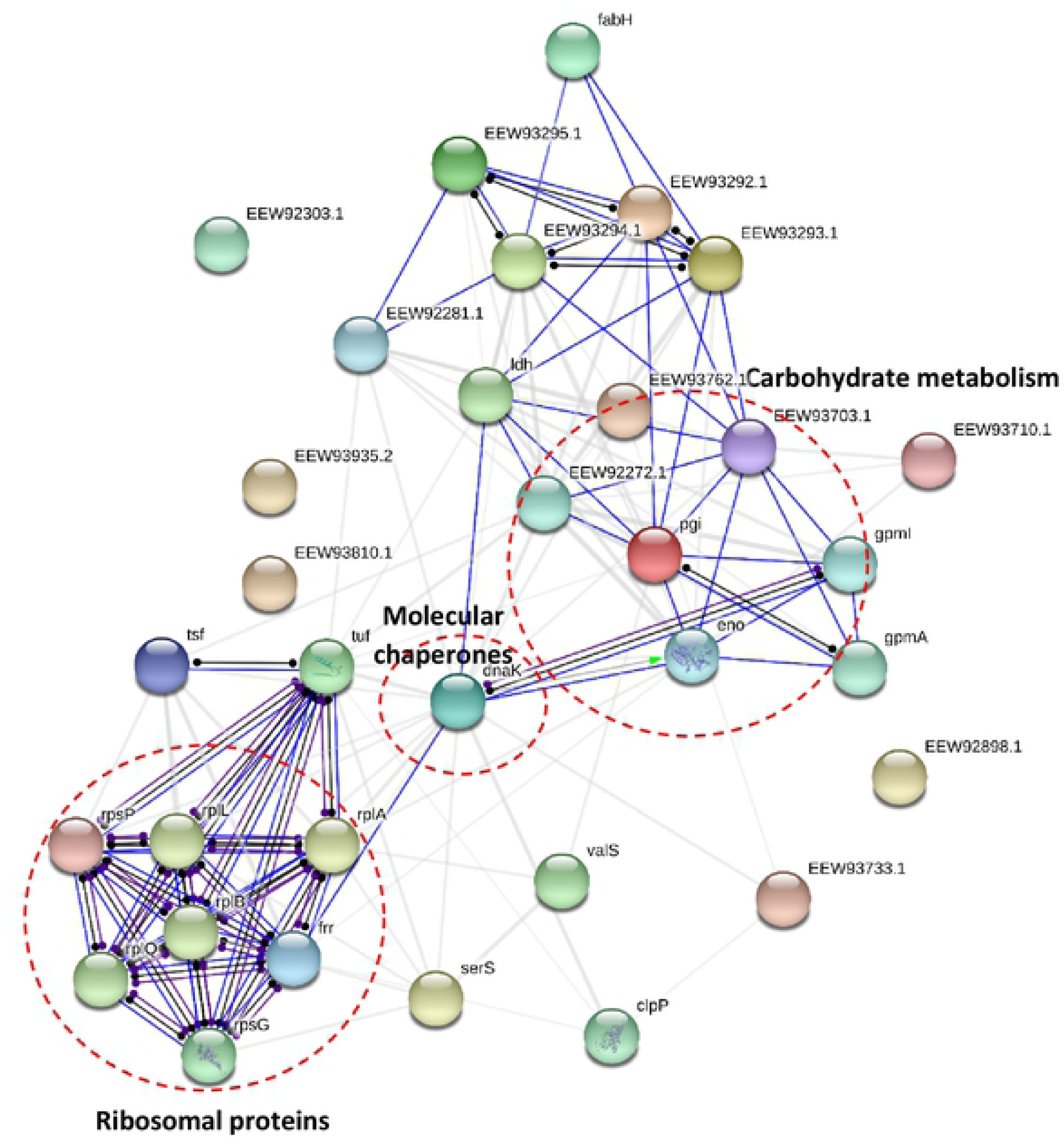
Functional protein association networks of *G. elegans* EVs proteome. The online tool STRING was used for grouping the EVs proteins based on functional networks. Minimum interaction scores were set at a strong confidence level of 0.7. The three major network groups formed are shown in dotted circles. Seven different colors link a number of nodes and represent seven types of evidence used in predicting associations. A red line indicates the presence of fusion evidence; a green line represents neighborhood evidence; a blue line represents co-occurrence evidence; a purple line represents experimental evidence; a yellow line represents text mining evidence; a light blue line represents database evidence and a black line represents co-expression evidence.

KEGG pathway analysis of the EVs proteomes is depicted in Fig 7. Proteins belonging to carbohydrate metabolism and genetic information processing were found to be the most predominant in *G. adiacens* and *G. elegans* EVs. About 37% of the proteins in *G. adiacens* EVs proteome was predicted to be involved in the carbohydrate metabolism and 25% in genetic information processing. On the contrary, *G. elegans* had the majority (37%) of the EVs proteins in the genetic information processing category followed by 23% in the carbohydrate metabolism category. As predicted by the pathway tool, a few proteins from both species were also implicated in amino acid metabolism, lipid metabolism, glycan metabolism, and energy metabolism. Vesicles equipped with metabolic machineries can help bacterial colonization and host cell invasion. For example, ATP generated in vesicles might regulate the activity of virulence factors and facilitate cell-cell communication of bacteria [49]. Overall, metabolism related proteins in the EVs might facilitate long-term contact between the bacterium and the epithelial cells, causing increased epithelial cell/tissue damage.

**Fig 7.**
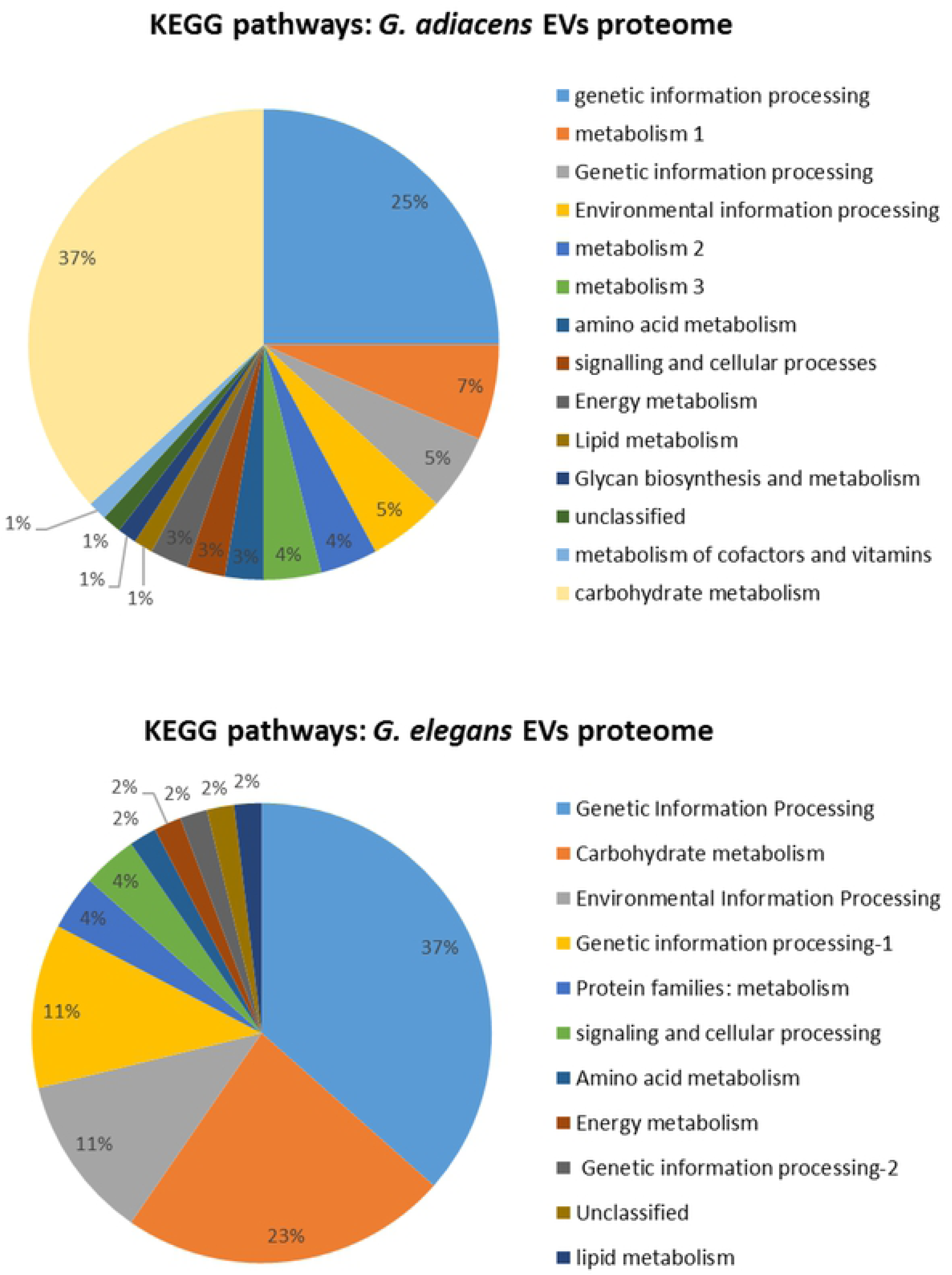
KEGG pathway analysis of the EVs proteomes. All protein sequences from *G. adiacens* and *G. elegans* vesicle proteomes were subject to KEGG pathway analysis using the genus “streptococcus” as reference.

#### Prediction of virulence proteins in the EVs proteomes

Our study revealed that EVs produced by both *Granulicatella* species contained proteins that were predicted to carry virulent properties. This finding overemphasizes the role of EVs in the pathogenesis of *Granulicatella* infections. Tables 2 and 3 show the list of 44 and 31 proteins that were predicted to be virulent from EVs proteomes of *G. adiacens* and *G. elegans*, respectively. The major proteins with demonstrated evidence on their virulence properties in other bacterial species were: thioredoxin [50], aminopeptidase [51], molecular chaperones DnaK and GroES [52, 53], Superoxide dismutase [54], Glyceraldehyde-3-phosphate dehydrogenase [55], phosphoglycerate kinase [56], and acyl carrier protein [57]. A vast literature on membrane vesicles has demonstrated that a number of well-known and extensively studied toxins and non-toxin virulence factors are secreted via vesicles [58]. Unlike virulence factors secreted in soluble form, vesicle-associated virulence factors are provided with a unique benefit of being protected from host proteases [13]. Moreover, vesicle-virulence factors are delivered to host cells/tissues as concentrated packages, increasing the damage level at specific target sites. Vesicle-mediated delivery of virulence factors is a widespread mechanism across bacterial species and genera. Similar to other oral bacteria such as *Aggregatibacter actinomycetemcomitans* [59], *Kingella kingae* [60] and others that are also implicated in infective endocarditis, *Granulicatella* species possibly use their EVs filled with numerous putative virulent proteins in the pathogenesis of this infection.

**Table 2.**
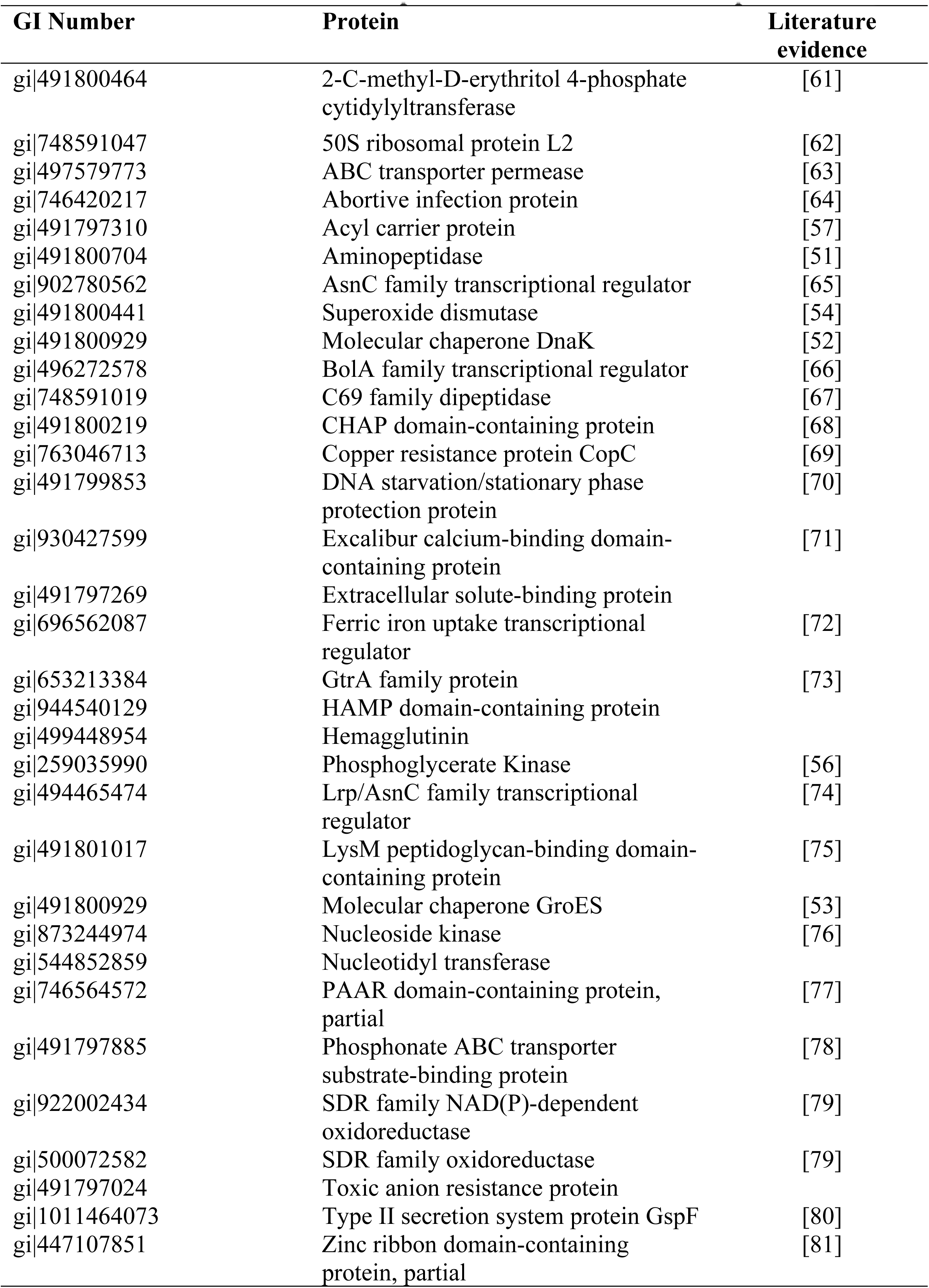
Putative virulence factors predicted in *G. adiacens* EVs proteome.

**Table 3.**
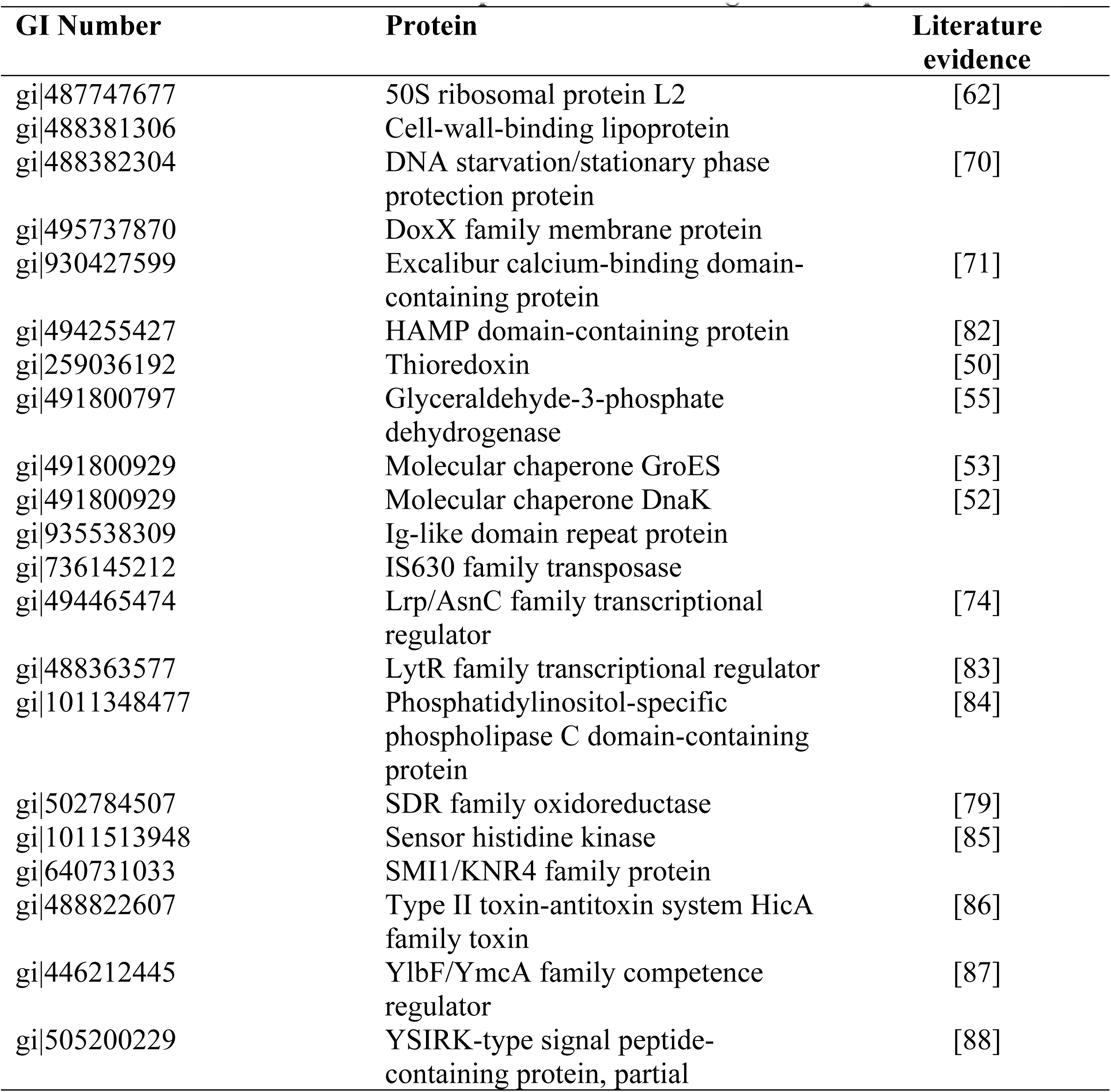
Putative virulence factors predicted in *G. elegans* EVs proteome.

#### ELISA quantification of selected cytokines produced from stimulated human PBMCs with EVs and WCP

As shown in Figures 8 and 9, all concentrations (10, 25, 50, and 100 µg/ml) of *G. adiacens* EVs, and *G. elegans* EVs triggered the production of the selected potent proinflammatory cytokines from human PBMCs as compared to the controls (0 µg/ml). Our study demonstrated that both *G. adiacens* EVs and *G. elegans* EVs were able to stimulate cytokine release from human PBMCs and thus could play a role in the induction of an inflammatory response. This finding is in accordance with previous studies [11, 14, 18, 89] that revealed the immuno-modulatory effects of EVs in other bacteria. In the current study, EVs from both species induced IL-8 and IL-1β, but not CCL5, in a dose-dependent manner. *G. adiacens* EVs induced the release of IL-8 and IL-1β to significantly (*P* < 0.05) higher levels compared to WCP (Fig 8). In the case of *G*. *elegans*, compared to WCP, EVs induced significantly higher levels of only the IL-1β (*P* < 0.05) (Fig 9). When cytokine levels were compared between the two species, no statistically significant difference was found. These observations overemphasize the importance of bacterial vesicle production in the activation of inflammation and thus pathogenesis of bacterial infections. The ability of bacterial vesicles to trigger host inflammatory response is a well-established phenomenon. When host epithelial cells encounter or take up the vesicles, an immediate innate immune response begins. IL-8 and IL-1β are prominent cytokines in infective endocarditis [90], but also in oral infections [91, 92]. IL-1β has a wide range of actions mediating inflammatory host response. At low concentrations, it mediates local inflammation while at high concentrations it possesses endocrine effects. Due to its neutrophil recruiting property, IL-8 is a major inflammatory cytokine induced by a variety of microbial components [93, 94].

**Fig 8.**
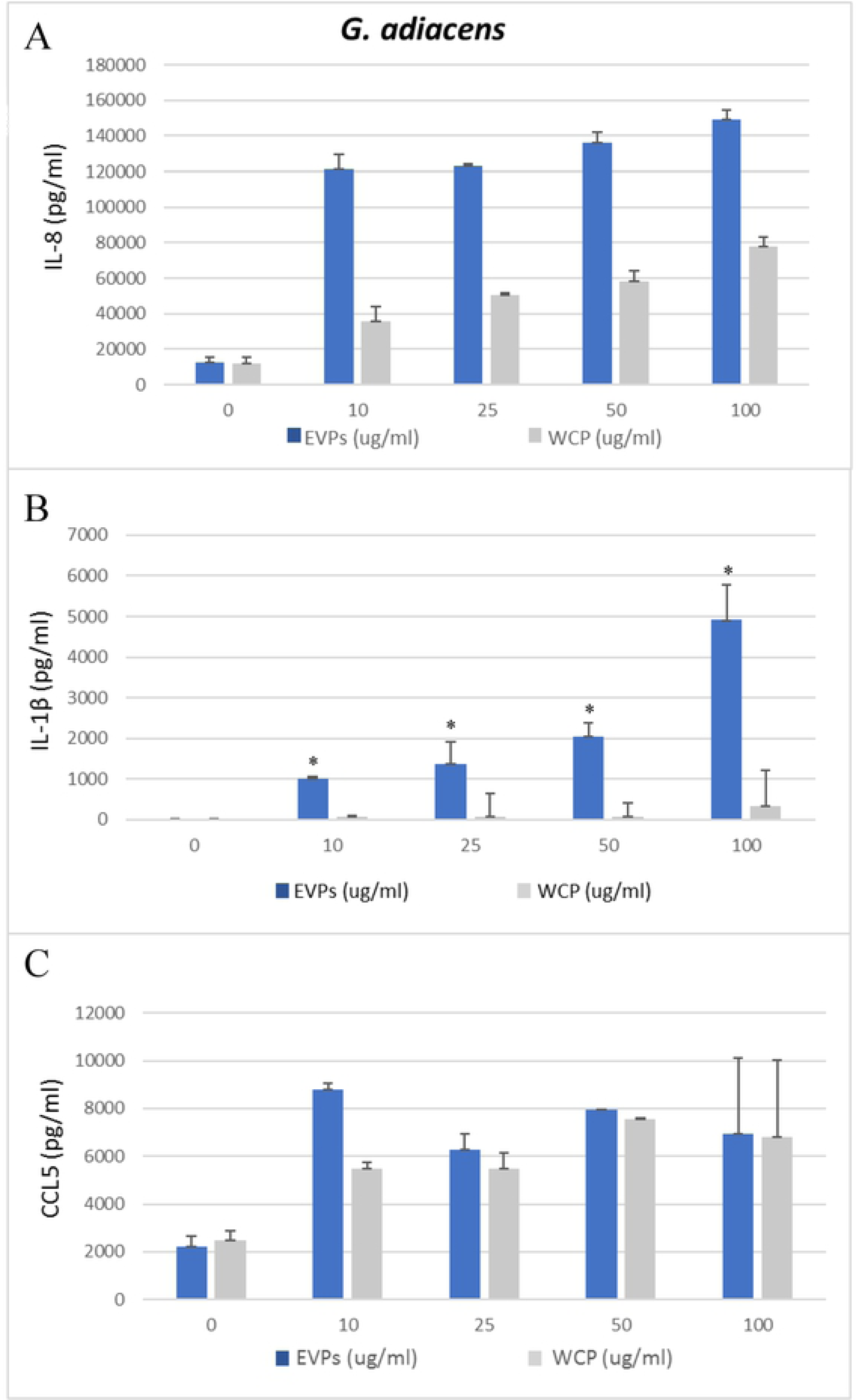
ELISA quantification of IL-8 (A), IL-1β (B), and CCL5 (C) production by human PBMCs stimulated with *G. adiacens* EVs and WCP. (10, 25, 50, and 100 µg/ml). EVs induction was considered significantly different from WCP induction at *p < 0.05.

**Fig 9.**
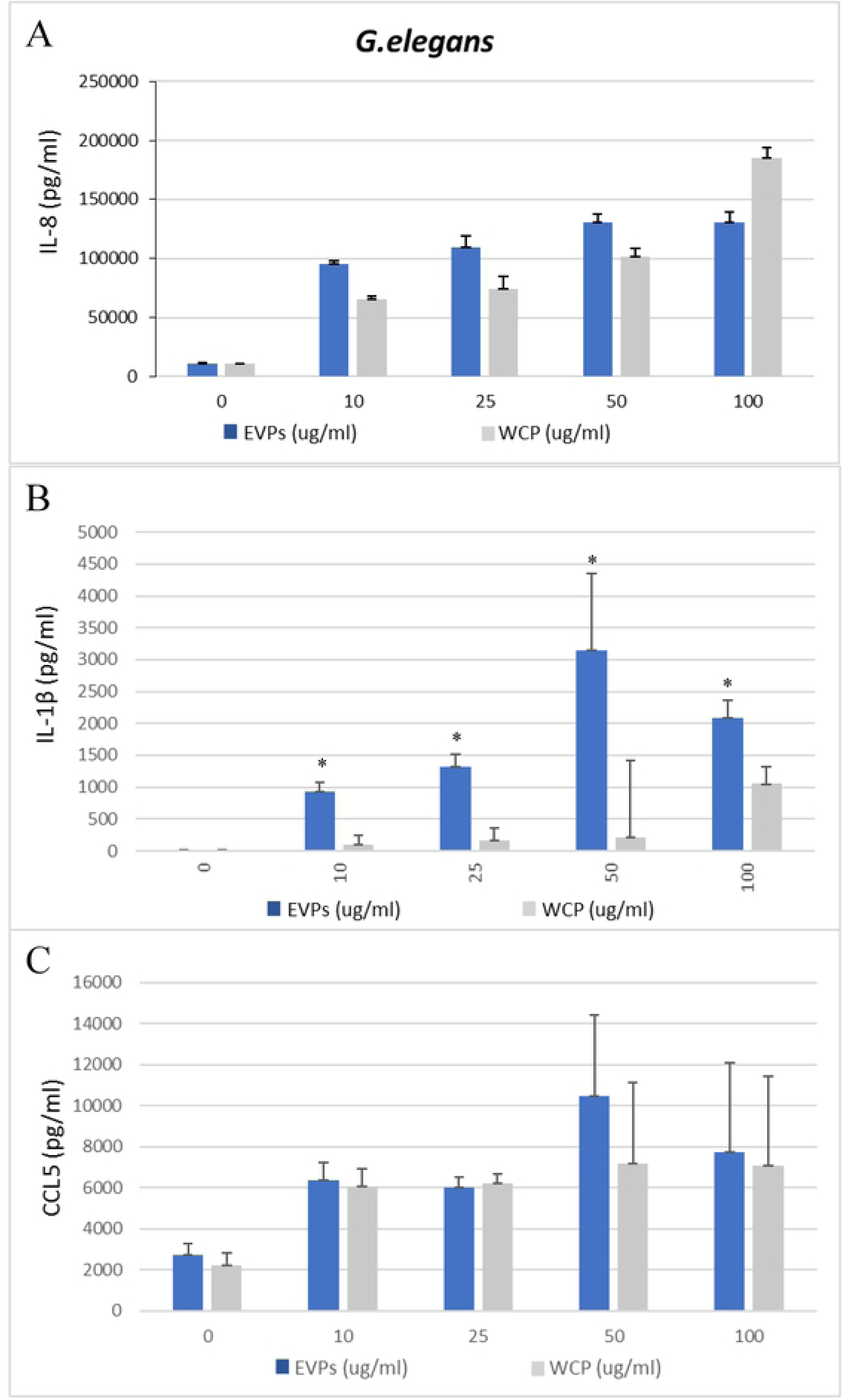
ELISA quantification of IL-8 (A), IL-1β (B), and CCL5 (C) production by human PBMCs stimulated with *G. elegans* EVs and WCP. (10, 25, 50, and 100 µg/ml). EVs induction was considered significantly different from WCP induction at *p < 0.05.

## Conclusion

To the best of our knowledge, this is the first research that presented evidence for the hypothesis that *Granulicatella* species release EVs. We discovered that the EVs proteomes of *G. adiacens* and *G. elegans* were enriched with a large number of predicted putative virulence factors. *Granulicatella* species possibly use their EVs as vehicles to deliver virulence factors to body sites not accessible to whole bacterial cells—a mechanism widespread across bacterial species. In addition to virulent proteins, which can impose direct detrimental effects on host cells/tissues, other components of the EVs, i.e., metabolic enzymes, ribosomal proteins, stress-response proteins may contribute to pathogenesis by enhancing adaptation of these species and survival in the hostile host environments. Thus, the diversity in EVs content emphasizes the possible roles of these vesicles in bacterial survival, invasion, host immune modulation as well as infection. Moreover, EVs of both species were demonstrated to be potent inducers of proinflammatory cytokines, and importantly, the EVs were significantly more potent than the whole cell proteins in eliciting inflammatory response. These EVs may play an important role in the activation of inflammation and thus pathogenesis of *Granulicatella* infections. Further functional characterization of the *Granulicatella* EVs may throw more light on how these species may utilize vesicles to orchestrate events that may lead them from being silent normal flora species towards infection-causing ones.

## Acknowledgments

This study was supported by Kuwait University Grant SRUL 01/14 and partly by the College of Graduate Studies, Kuwait University. Special gratitude is given to the Research Core Facility, Health Sciences Center, Kuwait University, the Department of Microbiology, Faculty of Medicine, Kuwait University, and The Nanoscopy Science Center, Faculty of Science, Kuwait University for the permission to use their facilities.

